# Retinotopic connectivity maps of human visual cortex with unconstrained eye movements

**DOI:** 10.1101/2023.03.16.533037

**Authors:** Gene T. Tangtartharakul, Catherine A. Morgan, Simon K. Rushton, D. Samuel Schwarzkopf

## Abstract

Human visual cortex contains topographic visual field maps whose organization can be revealed with retinotopic mapping. Unfortunately, constraints posed by standard mapping hinders its use in patients, atypical subject groups, and individuals at either end of the lifespan. This severely limits the conclusions we can draw about visual processing in such individuals. Here we present a novel data-driven method to estimate connective fields, fine-grained maps of the functional connectivity between brain areas. We find that inhibitory connectivity fields accompany, and often surround, facilitatory fields. The visual field extent of these inhibitory subfields falls off with cortical magnification. We further show that our method is robust to large eye movements and myopic defocus. Importantly, freed from the controlled stimulus conditions in standard mapping experiments, using entertaining stimuli and unconstrained eye movements our approach can generate retinotopic maps, including the periphery visual field hitherto only possible to map with special stimulus displays. Generally, our results show that the connective field method can gain knowledge about retinotopic architecture of visual cortex in patients and participants where this is at best difficult and confounded, if not impossible, with current methods.

## Introduction

The image of the world on the back of the eye is mapped onto the surface of brain. Adjacent neurons in the visual cortex code for responses of adjacent cells on the retina [Wandell et al., 2007]. This creates contiguous retinotopic maps in the visual cortex. Recent studies suggest that the human brain has over 50 retinotopic maps [Sereno et al., 2022; Wandell et al., 2007]. Functional magnetic resonance imaging (fMRI) is the primary method for exploring the maps in the human brain [Dumoulin and Wandell, 2008; Engel et al., 1997; Sereno et al., 1995]. Over a decade ago, the toolbox of visual neuroscience was extended with the population receptive field (pRF) method [Dumoulin and Wandell, 2008]. This method compares, voxel-by-voxel, the time course of changes in activity produced by the population of neurons contained within a voxel, to the time course of changes of visual stimulation within regions of the retinal image to determine the range of retinal locations that cause the voxel to respond. The result is a map of the “receptive fields” of each voxel. Use of the pRF method has effectively characterized organization of multiple human visual brain areas that code the parafoveal portion of space, permitting an understanding of their properties and stimulus preferences. However, this comes with several caveats: conventional pRF mapping varies the stimulus to systematically stimulate different parts of the retina. This stimulus-referred approach therefore demands steady fixation during the experiment. This is one reason why the representation of the foveal retina in visual cortex remains mostly unexplored. Moreover, pRF mapping also requires a clear and unimpeded view of the stimuli limiting the range of locations that can be stimulated. Therefore, the peripheral representation of the human visual cortex also remain mostly unexplored. Classical mapping experiments typically present stimuli via a screen mounted onto the head coil inside the scanner bore that only subtends the central 10-20° of the visual field. While some previous single-unit and fMRI studies attempted to minimize this limitation through use of eccentric fixation targets, goggles, telescopes, or spherical projection systems [Jolly et al., 2021; Mikellidou et al., 2017; Pitzalis et al., 2006; Smittenaar et al., 2016; Urale et al., 2022; Yu et al., 2012], the difficulty of such experiments, the requirement for special equipment, and the fact that most of these methods can still not stimulate the far periphery precludes us from studying peripheral visual field maps regularly in humans. This constrains our understanding of conditions affecting peripheral vision, such as glaucoma or hallucinations experienced in a range of disorders, like schizophrenia or neurodegenerative illness like Parkinson’s Disease because these typically occur in the peripheral visual field.

Connective field (CF) mapping has recently been proposed as an alternative way of investigating visual cortex architecture [Haak et al., 2012]. CF mapping relates the time course of activation of voxels in one brain region to the time course of activation of voxels in another region, usually, the primary visual cortex (V1. Thus, localized cortical activity in one region can effectively be modeled as a stimulus for neuronal populations in another region, shifting retinotopic mapping from stimulus- to neural-referred. This should theoretically obviate the requirement for steady fixation, and even for systematic visual stimuli, and could thus make retinotopic mapping more robust. Here, we test this prediction. We have developed a novel data-driven CF method that makes fewer assumptions than the forward-modeling procedure used previously [Gravel et al., 2014; Haak et al., 2012; Invernizzi et al., 2021; Knapen, 2021] and that is computationally highly efficient. We compare maps obtained with this method to those generated by conventional pRF analysis in the presence of large, erratic eye movements. Further, we evaluate CF maps obtained while participants play an entertaining video game without systematic stimulation or stable eye fixation.

## Materials and Methods

### Participants

In the *Standard Mapping* experiment, we collected data from 25 participants for a classical retinotopic mapping study (15 females; ages: 19-47 years). Two participants were excluded after retinotopic mapping analysis due to insufficient signal-to-noise ratios. In the *Unstable Eye* experiment, we collected data from 3 participants (2 females; ages: 20-31 years). In the *Laser Kiwi* experiment, we collected data from 5 participants (3 females; ages: 20-44 years). Two participants took part in both the *Standard Mapping* and *Unstable Eye* experiments. One of the authors took part in both the *Standard Mapping* and *Laser Kiwi* experiment.

In the *Standard Mapping* experiment, all participants had normal or corrected-to-normal visual acuity. In the *Unstable Eye* and *Laser Kiwi* experiments, three participants were myopic. For two of these, we deliberately left vision uncorrected because (1) they could not wear contact lenses and the eye tracker is unable to track reliably through spectacle lenses and (2) we were interested in testing the robustness of CF vs pRF analysis, respectively, to optical defocus. The remaining participants either had normal visual acuity or were corrected to normal by wearing their prescribed contact lenses. Participants had no other ocular pathologies. All participants were recruited from the local staff and student pool at the University of Auckland. They provided a written informed consent to participate in the study and the study procedures were approved by the University of Auckland Human Participants Ethics Committee (UAHPEC).

### Stimuli and task

The participants viewed the stimuli through the mirror subtending a visual angle of 31.8° horizontally and 17.8° vertically. The liquid crystal display screen (71 cm x 39 cm with 1920 × 1080 resolution and 120 Hz refresh rate; BOLDscreen, Cambridge Research Systems, Rochester, U.K.) displaying the stimuli was placed at the back of the scanner’s bore. Participants lay supine inside the bore. They viewed the screen through a mirror attached to the top of the head coil (total viewing distance in Standard Mapping experiment: 111cm; Experiments 2 and 3: 124.5cm. The curvature of the scanner bore blocked the view of the display’s top corners. Participants were instructed to keep still to minimize head motion. In Experiments 2 and 3, the participants’ eye movements were recorded with monocular gaze tracking using a magnetic resonance imaging-compatible eye tracker (EyeLink 1000+; SR Research, Ottawa, Canada), with data down-sampled to the video refresh rate of the screen. All stimuli were programmed in MATLAB R2021a (MathWorks, Natick, Massachusetts) using the Psychtoolbox 3 [Brainard, 1997; Pelli, 1997] (http://psychtoolbox.org).

#### Standard Mapping

We used our standard pRF mapping paradigm with sweeping bars as stimuli [Morgan and Schwarzkopf, 2019]. Specifically, a high-contrast ripple pattern [Schwarzkopf et al., 2014] was exposed through traversing bar apertures (width: 1°) on a uniform gray background. The bar traversed within a circular region with radius 9.5° centered on the screen. As the bar was restricted within this circular region, its length changed as it moved. The bar was presented at 4 different orientations, each with 2 sweep directions, producing 8 different configurations. Each run contained 8 sweeps of each bar configuration. The sweep occurred in the direction perpendicular to the bar orientation. Within a sweep, the bar traversed in 25 discrete steps, one step per TR (approximately 0.7° per second). The first sweep started from the horizontal orientation moving upward then changed by 45° clockwise each sweep. In between the fourth and fifth sweeps and after the last sweep, only the fixation dot was presented for 25s.

Participants were asked to fixate at a target dot (diameter: 0.09°) at the center of the screen. The scan duration was divided into epochs of 200ms. In each epoch, there was a 0.01 probability that the target dot changed color from blue to purple. The 200ms epoch immediately following a color change always only contained the fixation dot. Participants were asked to press a button on a magnetic-resonance compatible response box when they detected a color change. This acted as an attentional task, ensuring that the participant maintained their fixation on the dot. As in previous studies, we also included a low-contrast radar screen pattern [Morgan and Schwarzkopf, 2019] presented transparently on top of all the stimuli to maximize fixation stability.

The overall duration of a scanning run was 250 seconds (25s per 8 sweep directions and 2 blank epochs). All participants completed 6 runs of this experiment.

#### Unstable Eye

We adapted our standard pRF mapping paradigm to test the effect of unstable eye movements on retinotopic maps. A new scanner setup (see below) required a different viewing distance (see *Stimuli and task*); the radius of the stimulated part of the visual field was therefore only 9°. We also enlarged the fixation dot and enhanced its visibility if presented against the backdrop of the flickering stimuli (diameter: 0.33°, surrounded by a 0.17° width annulus of background gray). We further removed the radar screen pattern. Participants completed 8 runs but we divided the runs into two types – stable and random. In the stable runs, the dot was always positioned centrally (Figure 1A, left), whereas in the random runs, the dot had a 0.003 probability on each screen refresh (120Hz; on average it would move once every 2.8s) of switching to a new random location within a set square region (Figure 1A, middle). This square was centered on the screen and occupied 30% of the screen height. The dot appeared blue if the participant fixated at the location. However, if the gaze position was recorded more than 1.7° from the dot center, the dot color changed to red. This encouraged the participant to chase the dot with their gaze. In addition, they also performed a task on the fixation dot. The scan duration was divided into epochs of 200ms. In each epoch, there was a 0.01 probability that the target dot changed into either a Latin letter or a single-digit number. Participants were instructed to respond with a button press whenever a number appeared.

**Figure 1.**
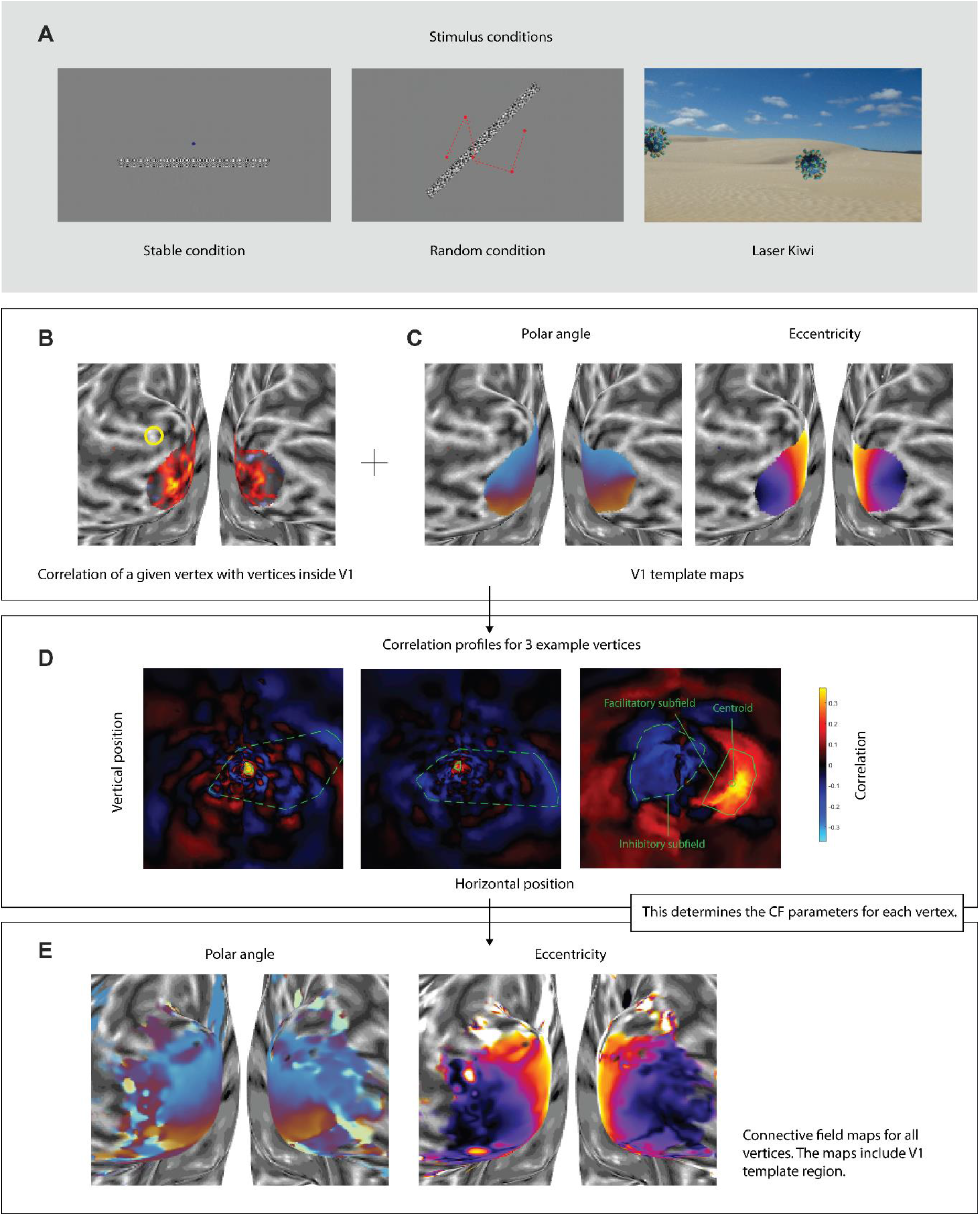
Connective field analysis based on reverse correlation. (**A**) Stimulus examples from the stable and random conditions in *Unstable Eye* experiment (left and middle, respectively), and an example frame from the *Laser Kiwi* experiment (right). Note that the stimulus for *Standard Mapping* is not shown here, but this was similar to the stable condition (see *Materials and Methods* for details). (**B**) Correlation between the time series for a particular vertex (denoted by yellow circle) and the time series of vertices inside V1 delineated using Benson probabilistic atlas. Hotter colors indicate stronger correlations. (**C**) V1 polar angle and eccentricity map derived from Benson probabilistic retinotopic map used as templates. (**D**) Correlation profiles of three example vertices projected back into the visual field using the template map. Each profile shows the centroid of the CF (green circle), facilitatory (solid line) and inhibitory connective subfields (dashed green line). (**E**) Polar and eccentricity map of CFs estimated for all vertices in the occipital cortex (including vertices inside the template region V1 to estimate the correlations of template vertices with its neighbors). Data in **B, C**, and **E** are shown on an inflated model of the gray-white matter boundary. Plots in **D** are in visual space.

#### Laser Kiwi

Participants were presented with a gaze-contingent interactive game inspired by a popular entry to the 2015 New Zealand flag referendum [Gray, 2015]. The game displayed coronavirus images (https://commons.wikimedia.org/wiki/File:Coronavirus._SARS-CoV-2.png) floating across the screen (Figure 1A, right). During the runs, the participants were asked to role-play as a superhero shooting laser eyes at the targets. A green dot on the screen (diameter: 0.33°, with a smoothed edge where transparency ramped up), showed the participant’s eye position. The viruses had a radius of 4.6°. At every screen refresh, there was a 0.005 probability of spawning a new virus (on average once every 1.7s) anywhere on the screen with a maximum of 6 viruses appearing simultaneously. The viruses traveled at a speed of 0.08° (2 pixels) per screen refresh (approximately 10°/s) in a random linear direction and rotated (randomized clockwise or counterclockwise) by 1° per screen refresh.

The participants needed to fixate within 2° from the center of a virus to eliminate it. Viruses appeared with varying hit points, randomized between 20-180 determining for how long they must be fixated to be eliminated. For every screen refresh that the eye gaze was recorded as hitting the target, a hit point was subtracted. Whenever a virus had fewer than 20 hit points, it started to shrink and jitter by adding a random orientation at every screen refresh. When it had zero hit points, we displayed an explosion effect of expanding white discs with increasing transparency and played a corresponding sound effect.

The viruses were overlaid on top of background images depicting outdoor scenes, buildings, people, animals, writing in various scripts, and textures. Each background was shown for 5 seconds in a shuffled order, providing a total of 72 backgrounds per run. The images filled the whole screen and therefore subtended a visual angle of 32° horizontally and 18° vertically.

### MRI data acquisition

Data were collected at the University of Auckland’s Center for Advanced Magnetic Resonance Imaging (CAMRI). In the *Standard Mapping* experiment, we acquired data with a Siemens MAGNETOM Skyra 3 Tesla scanner. Following a scanner upgrade in 2022, data from the *Unstable Eye* and *Laser Kiwi* experiments were acquired with a Siemens MAGNETOM Vida Fit 3 Tesla scanner. In all functional experiments, the same 32-channel head coil was used, with the front removed to allow an unimpeded view of the screen resulting in remaining 20 effective channels covering the side and the back of the head.

We collected participants’ functional data using T2*-weighted echo-planar imaging. Scan parameters were matched before and after the upgrade using an accelerated multiband sequence (2.3 mm isotropic voxel resolution, 96 × 96 matrix size, 62° flip angle, repetition time (TR): 1000 ms, echo time (TE): 30 ms) with 36 transverse slices angled to be approximately parallel to the calcarine sulcus. The scan had a multiband/slice acceleration factor of 3, an in-plane/parallel imagine acceleration factor of 2, and rBW was 1680 Hz/Px.

The number of volumes and runs varied across the experiments. In the *Standard Mapping* and *Unstable Eye* experiments, we collected runs comprising 250 T2*-weighted image volumes each. In the *Standard Mapping* experiment, we collected 6 such runs per participant. In the *Unstable Eye* experiment, we collected 8 runs per participant (alternating runs contained the stable and random conditions, respectively). In the *Laser Kiwi* experiment, we collected 2-4 runs, comprising 360 volumes per run.

We also acquired the participant’s T1-weighted structural image for the coregistration of functional and structural images using magnetization-prepared rapid acquisition with gradient echo scan (MPRAGE; collected with 1 mm isotropic voxel size, 8° flip angle, 880 ms inversion time (TI), 2000ms TR, 2.8ms TE, and 208 sagittal slices for full brain coverage, taking 4 min 56 s). The front of the head coil was put back on to collect this structural scan, to improve signal to noise ratio).

### Preprocessing

We used the default parameters of SPM12 (Wellcome Centre for Human Neuroimaging, London, UK) to realign and co-register our functional data to the structural scan for each participant. Surface mesh models of the gray-white matter and pial boundaries were reconstructed and inflated with the automatic reconstruction algorithm in FreeSurfer (7.1.1; https://surfer.nmr.mgh.harvard.edu). We then projected the functional data onto the cortical surface model. To this end, we used the *mri_vol2surf* function in FreeSurfer to determine the voxel in the functional image that locates halfway between the gray-white matter and pial boundaries for each vertex on the surface mesh. During this stage, data were smoothed along the surface with a kernel of 3mm. This reduces partial volume effects and fills in signal loss that would otherwise arise in a nearest neighbor interpolation of voxels to the surface mesh.

All further analyses were conducted using SamSrf 9 (https://osf.io/2rgsm). For each vertex, we applied linear detrending to their time series, removing the slow drifts, and subsequently normalized them to z-scores. To save time and computing resources we only analyzed data from an occipital region of interest by selecting vertices in the inflated surface model whose Y-coordinates (anterior-posterior axis) were below -35.

### Connective field analysis

Each vertex in the brain can have a connective field in V1. This corresponds to the portion of V1 that is functionally connected to that vertex, that is, where V1 responses coincide with its own activation. Previous implementations of connective field analysis [Haak et al., 2012] characterize the CF in terms of its anatomical location and extent within V1. This provides information about how functional connectivity is organized in cortical space. CF size can be quantified in terms of its geodesic extent across the cortical surface. However, this has some significant caveats, most notably the fact that the template region, V1, is split across the two cortical hemispheres. This creates an ambiguity when a CF is near the upper or lower border of V1, corresponding to the vertical visual field meridian. Most previous CF studies simply restricted the analysis to separate cortical hemispheres, but that loses information about whether a given CF spans the vertical meridian. It is also impossible to quantify the size of such a CF in terms of geodesic distance. Another caveat is that previous CF procedures tacitly assume the CF to conform to a particular shape, usually a two-dimensional Gaussian on the cortical surface. It remains unclear how appropriate this assumption is.

Crucially, because V1 contains a retinotopic map of the visual field this CF also corresponds to a portion of the visual field (Figure 1B,C). Thus, it is possible to estimate the visual field location of any given CF by determining the visual field location it represents within the V1 map. This circumvents the issues with CFs spanning the anatomical discontinuity of the vertical meridian. To this end, we used a probabilistic atlas prediction [Benson et al., 2012] to create a template retinotopic map of V1. We then used this V1 region as our template region for connective field modeling.

We assume that a neuronal population outside of V1 with a similar receptive field to a collection of vertices in V1 will elicit a similar time course of activation. We therefore calculated the correlation between the time series of a particular vertex with the time course of each vertex in V1 (Figure 1B). Note that this correlation can be either positive or negative. Next, using the probabilistic template retinotopic map (Figure 1C), we mapped the correlation profile within V1 back into visual space (Figure 1D). Instead of fitting a model to estimate CF shape, we used a data-driven estimation of its position and extent. Using a region-growing approach, we selected the contiguous region around the peak correlation where correlations were above half-maximum. We determined the outline of this profile (green solid line) using a convex hull algorithm, reflecting the extent of the facilitatory connective field. We used the convex hull centroid to estimate the visual field location of the CF (green circle), and the square root of its area to quantify its size. Additionally, we estimated the size of the inhibitory CF by finding the peak negative correlation, selecting visual field locations (again using a convex hull algorithm) where correlations were below the half-peak negative correlation (dashed green line), and then calculating the square root of this area. All these measures are therefore in visual space. Finally, we used the peak negative correlation to quantify the overall strength of inhibition. These CF parameters, centroid location, facilitatory and inhibitory size, as well as suppression strength, can then be plotted on the cortical maps to characterize retinotopic architecture similar to how pRF parameters are plotted on the cortical surface in conventional mapping (Figure 1E, for polar and eccentricity maps based on CF estimates).

### Population receptive field analysis

We also conducted a standard forward-modeling pRF analysis [Dumoulin and Wandell, 2008; Moutsiana et al., 2016; Urale et al., 2022] on the data from the *Standard Mapping* and *Unstable Eye* experiments. This was to compare our CF estimates to conventional pRF measurements and to evaluate the impact erratic eye-movements have on retinotopic mapping with either method.

We characterized the pRF for each vertex on the cortical surface. We assumed each vertex contains a population of neurons with receptive fields that respond to stimuli at a similar retinotopic location. We modeled pRFs as a two-dimensional Gaussian quantified by three main parameters: retinal position (x,y) and pRF size (σ). To achieve this, we used a coarse-to-fine approach.

We first performed an extensive grid search to find the best-fitting model by generating thousands of candidate pRFs that differed in pRF location parameters x, y and pRF size, σ. The moving bar aperture revealing the stimulus was defined as a binary mask, on a 100×100 grid of visual field locations. We predicted the response of each candidate pRF from the overlap of the pRF and the stimulus aperture. The response was then convolved with a canonical hemodynamic response function [de Haas et al., 2014] to account for the delay in the blood oxygenation-dependent response. For each vertex, the candidate pRF with the highest correlation between the predicted time course and the measured time course was selected.

The set of parameters of the candidate pRF with the highest correlation was then used in an optimization algorithm to estimate the pRF parameters more precisely. Only vertices with R^2^>0.01 in the grid search were included in this optimization stage. Vertices with x, y, or σ parameters greater/less than twice the size of the stimulated portion of the visual field were excluded as artifactual estimates. Because σ is sign-invariant and the optimization procedure does not constrain parameter estimates, we subsequently took the absolute value of σ as an estimate of pRF size.

### Regions of interest (ROI)

In the *Standard Mapping* experiment, we manually delineated visual cortical regions V1, V2, V3, V3A, V3B, and V4 based on the polar angle reversals [Sereno et al., 1995; Wandell et al., 2007] and the extent of activation in the pRF maps with a goodness-of-fit with p<0.0001. Our purpose in this experiment was to compare the pRF maps to maps from our novel CF method. Therefore, we wanted an accurate description of the map architecture as derived from conventional analysis.

In Experiments 2 and 3 we chose a different approach. The CF maps from the *Standard Mapping* experiment revealed significant responses for retinal locations peripheral to the coverage of conventionally analyzed pRF maps. This was expected because the CF method does not strictly require a visual stimulus and can theoretically even work with resting-state data [Gravel et al., 2014; Invernizzi et al., 2021; Knapen, 2021]. Restricting our analysis to the regions that can be defined based on the pRF maps would therefore exclude these additional visual field maps. Moreover, since we were interested in comparing the clarity and stability of retinotopic maps between the two methods, basing the delineation on maps obtained with one method would potentially bias the results. For our quantitative analyses we therefore used a probabilistic atlas generated by neighbor-preserving topological maps [Sereno et al., 2022], to define regions of interest. Using a nearest neighbor algorithm, we warped this atlas from the *fsaverage* template brain space into each participant’s native brain space. Such atlas maps inevitably fail to entirely accurately capture all the peculiarities of an individual’s retinotopic architecture due to individual differences in brain architecture and/or there are errors with spatial normalization. However, the delineation is fully reproducible and free of any experimenter bias. We used the delineations for estimating summary statistics of pRF and CF properties. Due to the atlas inaccuracies these summary statistics may therefore combine vertices that cross regional boundaries in some cases and should not be taken as a perfect reflection of a given brain region. Rather, they quantify consistent trends related to anatomical landmarks.

Using the atlas maps, we investigated all early visual areas from V1-V4, and V3A/B. Since we were also interested in regions representing the peripheral visual field, we further included V6, medial temporal (MT) areas, area prostriata (ProSt) and early intraparietal sulcus (IPS). Since the Sereno atlas defines numerous small areas, we combined some regions to create larger clusters. Our MT ROI consisted of upper and lower middle temporal regions (MT-, MT+), middle crescent (MTc, equivalent of V4t), and ventral and dorsal medial superior temporal regions. Our ProSt ROI included areas prostriata 1 and 2, while our IPS ROI included V7, lateral intraparietal areas (LIP0, LIP1), caudal intraparietal sulcus (cIPS), and caudal parietal area E (PEc). However, MT and IPS regions defined this way contained insufficient CF data and we therefore excluded these regions from our quantitative analysis.

## Results

We developed a novel data-driven procedure for estimating connective fields from fMRI data. This method makes fewer assumptions than previous CF implementations [Haak et al., 2012; Invernizzi et al., 2022; Knapen, 2021] because it simply characterizes the location and extent of CFs in terms of visual space, rather than fitting a formal model. This method also circumvents issues with anatomical separation of the two cortical hemispheres because CFs can span the vertical meridian (corresponding to the hemispheric separation between left and right V1).

### Standard Mapping experiment

To compare our new CF method to conventional pRF maps, we used a data set obtained using a standard pRF mapping protocol. CF maps reproduced the typical retinotopic architecture in occipital cortex (Figure 2). The contrast of CF maps was greater, with more pronounced polar angle reversals and eccentricity gradients than in standard pRF maps. However, the gradients, especially for polar angle maps, were also notably coarser and somewhat “noisier” in CF maps, and some polar angle reversals were broken up by ipsilateral position estimates or other likely artifacts. Nevertheless, the functional architecture of visual regions V1-V3 was clearly identifiable, and in many participants V4 and V3A/B could also be easily differentiated. Interestingly, conventional pRF maps often contained a lot of errant (presumably artifactual) pRF estimates outside the delineated visual regions. We rarely observed this in the CF maps.

**Figure 2.**
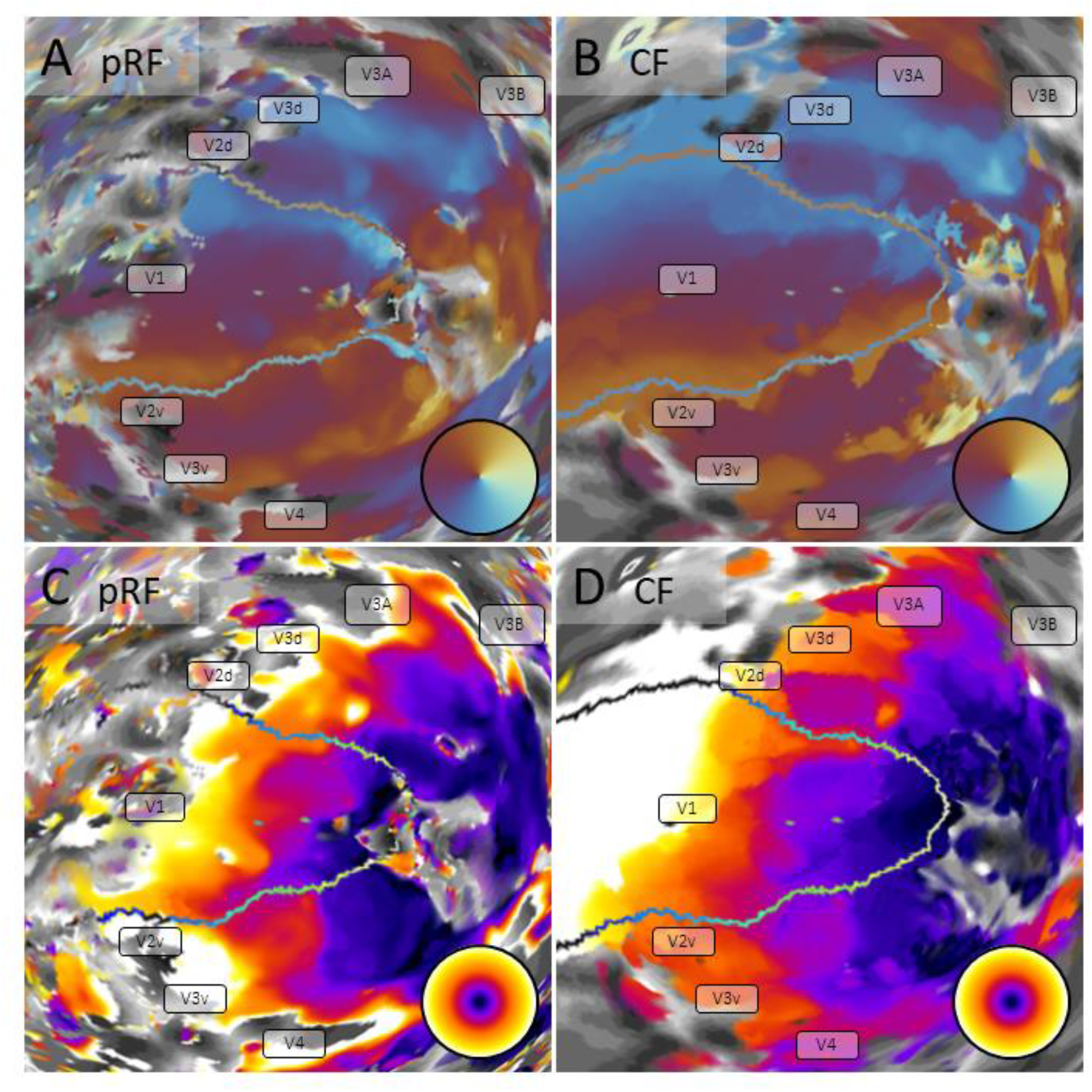
Retinotopic maps in a representative participant derived from conventional pRF mapping (**A**,**C**) compared to CF mapping (**B**,**D**). Maps for polar angle (**A**,**B**) and eccentricity (**C**,**D**) (see color wheel insets) are shown on an inflated spherical model of the left cortical hemisphere. Greyscale patterns indicate the cortical folding pattern. The transparent outline shows the borders of the template region V1.

Note that because we used a template map of V1 as template region for the functional connectivity analysis, the CF maps cover this atlas V1 in its entirety; analysis of this region is necessarily circular because each vertex in the template region must be perfectly correlated with itself. We nevertheless decided to retain this part of the map for two reasons. First, this allows us to quantify the similarity between this template-based CF map of V1 and the empirical pRF maps. Second, this also yields an estimate of the local connective fields within the template region, that is, how responses of a given vertex correlate with those of its cortical neighbors. Outside V1, however, we also note that CF maps (Figure 2B,D) extended considerably farther into the representation of the visual periphery than the eccentricity of the mapping stimulus. In contrast, in pRF maps (Figure 2A,C) visual responses ceased abruptly beyond about 12° eccentricity.

Next, we quantified parameter estimates for these maps across the whole sample. First, to visualize how CF sizes change across the visual field, we plotted vertex-wise CF sizes against their corresponding eccentricity estimates, pooled across all participants (Figure 3, black density histograms). To quantify this, we then fit a robust linear regression to those data. These regressions were conducted separately for each participant, and the coefficients then averaged across the group. To compare this with conventional pRF mapping, we conducted the same analysis for pRF sizes.

**Figure 3.**
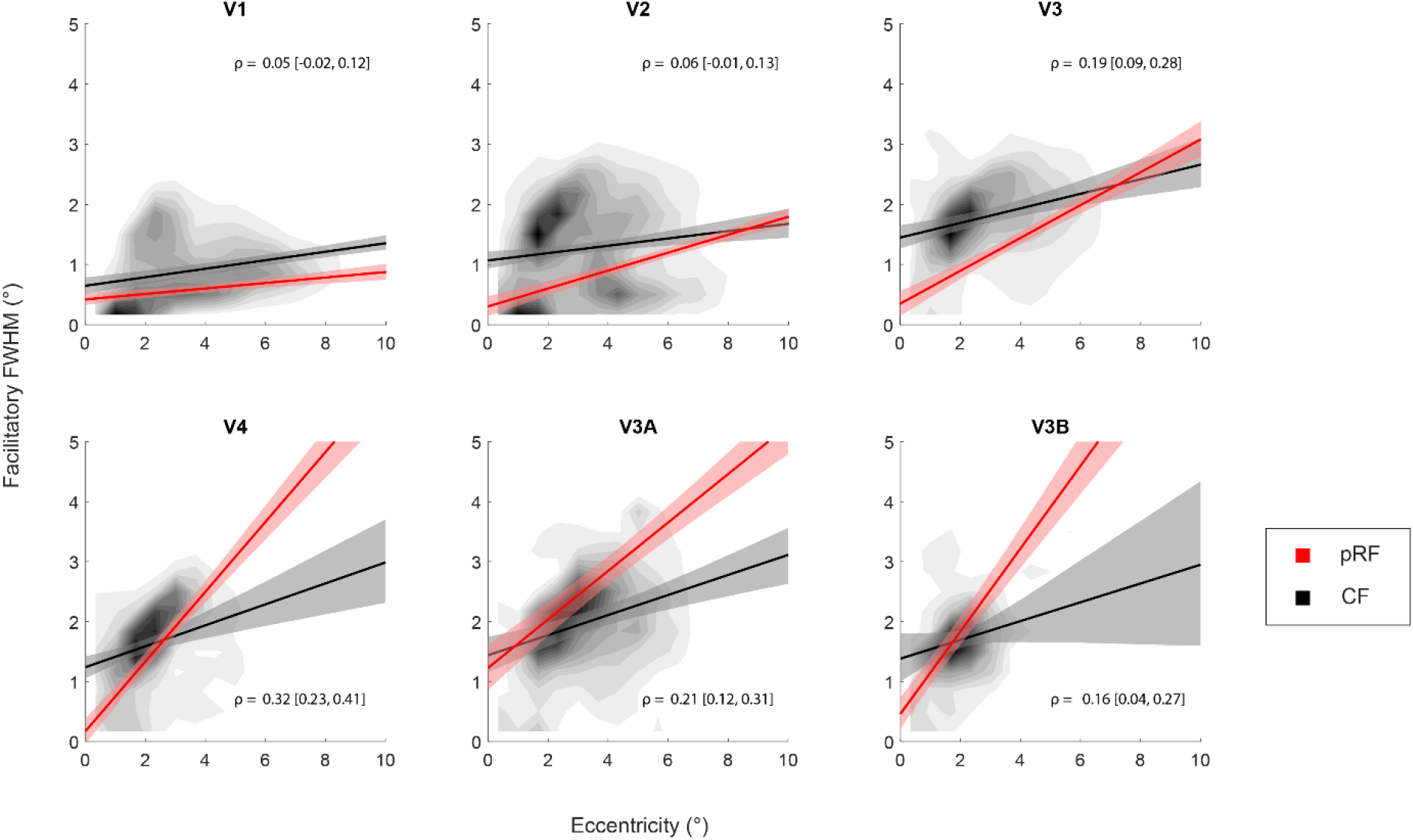
Comparison of CF sizes and pRF sizes in visual space. Each panel shows data from one visual region of interest. The solid lines show the average linear regression of CF (black) and pRF (red) size, respectively, as a function of eccentricity. Shaded regions denote the bootstrapped 95% confidence interval of the mean across participants. The two-dimensional density histogram visualizes CF sizes and eccentricities of all individual vertices (pooled across all participants). The statistics in each panel are the mean Spearman correlation (and bootstrapped 95% confidence interval) across participants between individual CF and pRF size estimates.

For both methods, CF/pRF sizes increased significantly with eccentricity (Figure 3, black and red curves), but the relationship between CF size and eccentricity is different than for pRF sizes. The slopes become increasingly steeper for pRF sizes in higher visual regions than in V1. In contrast, while the slopes for CF sizes also increased subtly across the visual hierarchy, they were much shallower than for pRFs. Slopes were also similar for V3, V4, and V3A/B. To further determine how comparable the spatial patterns of these estimates were, we calculated the Spearman correlation between pRF and CF size estimates. These correlations were also calculated in each participant and then averaged across the group. We bootstrapped these average correlations by resampling the values with replacement 10,000 times. A correlation was considered significant, if the 95% confidence interval does not overlap zero. This showed that pRF and CF size were significantly correlated only in V3, V4, and V3A/B (Figure 3, statistics shown in each panel).

In addition to the facilitatory CF size, our method also estimated the inhibitory CF profile where responses in a target vertex were inversely correlated with the vertices in the template region. This could reflect true inhibitory neural processing or could be a consequences of blood stealing by activated neuronal population. Inspection of individual CF profiles projected in visual space revealed different shapes of this inhibitory subfield (Figure 1C). In some target vertices, the inhibitory subfield was very pronounced and surrounded the facilitatory subfield. These inhibitory surrounds were typically very large, extending well into the peripheral visual field. However, in other CF profiles, the inhibitory subfields were shifted relative to the facilitatory region, and in some cases entirely spatially separated. Both types of inhibitory subfields often crossed the vertical meridian, corresponding to the hemispheric separation of primary visual cortex. Finally, some CFs showed only weak evidence of any inhibitory subfields.

We then quantified the inhibitory CF parameters, specifically the extent of the subfield and the peak negative correlation. The size of the inhibitory subfield (Fig 4A) suggests an inverse relationship with eccentricity, with large sizes in the parafovea below ∼2° eccentricity but then exponentially decreasing towards the peripheral visual field (the exponential relationship is most noticeable in V3 and V3A where there is a gentle bend between 2-3°). We again fit a robust linear regression to the vertex-wise data in each participant and visual region after logarithmic transformation of the inhibitory CF size estimates and then averaged the regression coefficients across the group. This demonstrated that in all regions the size of inhibitory CF subfields decreased significantly with eccentricity (Supplementary Figure S1A,B). We also conducted a similar analysis for the strength of inhibition, the peak negative correlation across the CF profile (Figure 4B). This showed that inhibition was significantly below zero but tended to increase with eccentricity (Supplementary Figure S1C,D), except in V3B, possibly because of the narrow range of eccentricities in that region.

**Figure 4.**
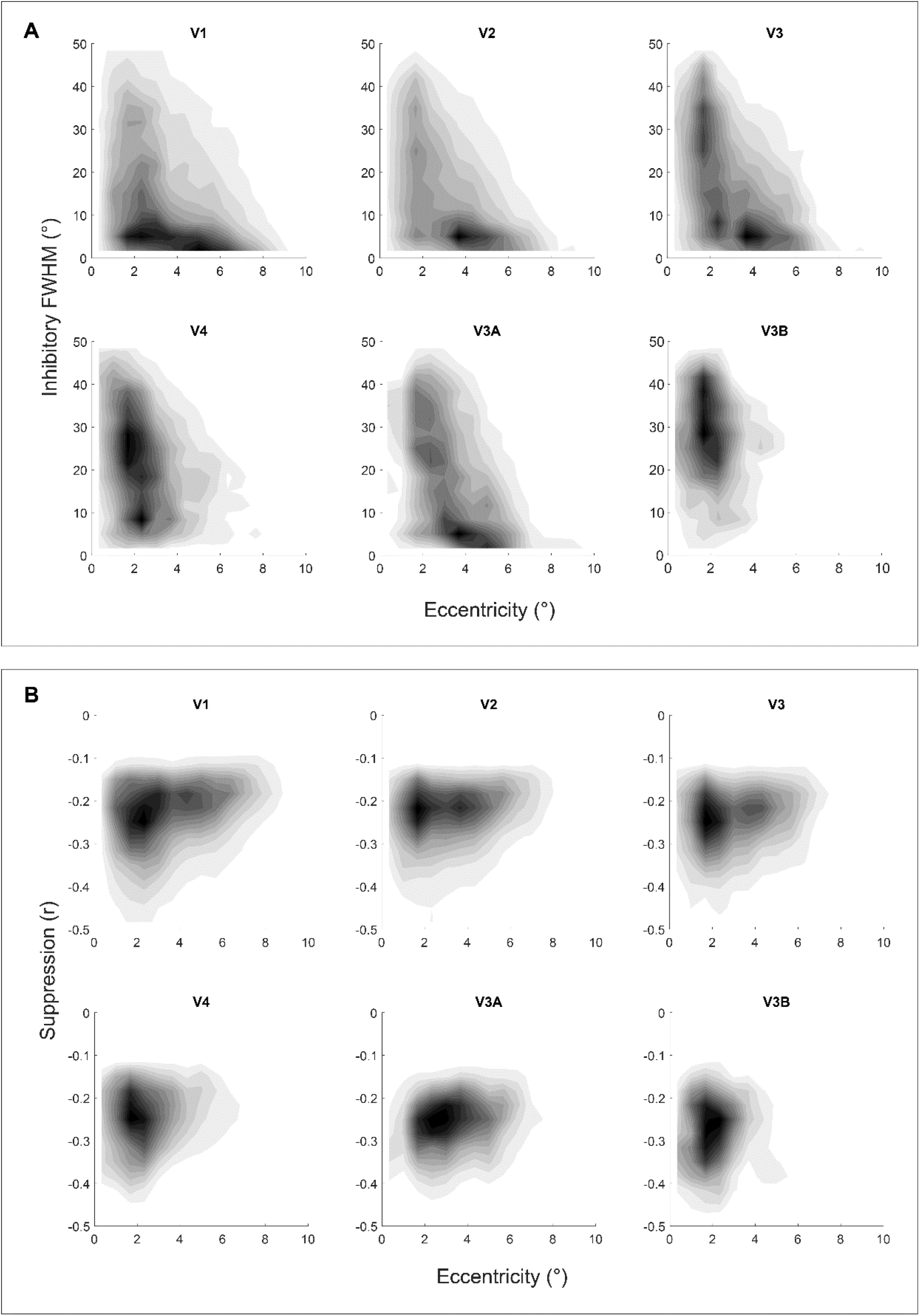
Parameters of inhibitory CF subregions. The two-dimensional density histograms plot eccentricity for individual vertices against inhibitory CF size (**A**) and suppression strength (**B**). Data were pooled across all participants.

### Unstable Eye experiment

Having implemented the novel CF method, we next aimed to investigate the theoretical robustness of the CF method against eye-movements. We again used a standard pRF mapping paradigm with high-contrast bars traversing the visual field. Participants were asked to have steady fixation (stable condition; Figure 1A, left) or to follow the fixation dot with their gaze while it jumped randomly (random condition; Figure 1A, middle). Using conventional pRF mapping, the presence of eye movements should add considerable variability to the retinotopic maps. Eye-movements will displace the retinal image. The stimulus-referred nature of the analysis means that unsteady eye fixation therefore breaks the correspondence between the assumed stimulus model and the actual retinal field locations stimulated at any given time. As a result, this should distort the retinotopic maps and reduce visual field coverage through data loss. It might also increase overall pRF size estimates because a wider range of locations is being stimulated for each vertex than under stable fixation. Our random condition was designed to induce such movements. We hypothesized that this would significantly impact the quality of the maps generated by the pRF but not the CF method, because the latter are based on cortico-cortical connectivity rather than the retinal projection to the cortex.

Side-by-side comparison of retinotopic maps for one example participant (Figure 5) from both stable and random conditions suggests that the pRF maps contained less activation throughout the visual regions due to eye-movements. In the polar angle map (Figure 5A), some of the expected retinotopic representation remained in early regions V1, V2, V3, and V3A, but this was incomplete, especially around the foveal representation, and the polar angle gradients were somewhat erratic. Especially on the lateral side of the occipital pole the organization also changed dramatically from the lower to the upper vertical meridian. pRF analysis provided no information for the higher atlas regions of V4, V6, MT+, or IPS. Eccentricity representations in the random condition of the pRF map were also erratic and lacked a clear peripheral representation (Figure 5B). This participant in fact showed the clearest pRF maps in the presence of eye-movements. In the other participants, little structure could be discerned in pRF maps obtained with unstable fixation (Supplementary Figure S2).

**Figure 5.**
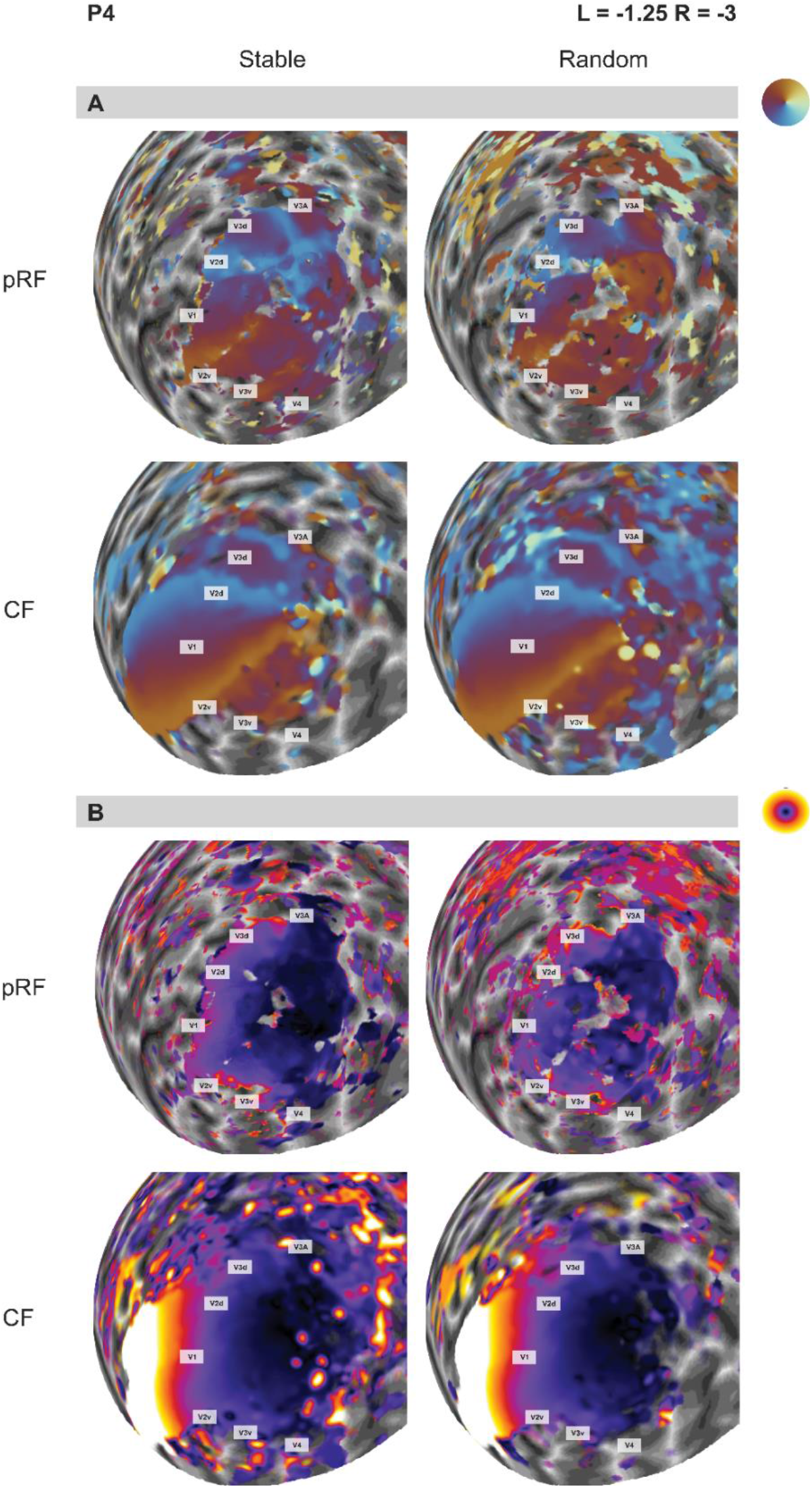
The effect of large eye-movements on retinotopic maps in the *Unstable Eye* experiment. Polar angle (**A**) and eccentricity maps (**B**) of participant P4 were derived by pRF or CF analysis (rows), either with stable or random fixation patterns (columns). This participant showed the clearest pRF maps with unconstrained eye-movements. See Supplementary Figure S2 for maps from all participants.

In contrast, CF maps were largely unaffected by these eye movements (Figure 5). Note again that being the template region, V1 of course contained a full contralateral visual field representation. However, even beyond V1 the CF maps showed wide-spread activation and with the expected retinotopic organization in both experimental conditions. Reversals at the vertical meridian indicating regional borders were largely consistent with the pRF maps from the stable condition. Ipsilateral visual field representations were rare in the CF maps. Eccentricity maps also showed the expected gradient between central and peripheral representations (Figure 5B). The eccentricity increased along the posterior-anterior axis. The foveal confluence of V1, V2, V3, and V4 could also be distinguished.

The task in the stable condition was a standard mapping paradigm. pRF maps generated without eye movements were therefore far less patchy and variable, and contained more widespread activation than in the random condition. Nevertheless, in comparison to the CF maps, both the overall extent of activation was substantially reduced. Within the part of cortex corresponding to the visual stimulus, the retinotopic architecture was very similar between the two analysis methods. This suggests that CF modeling was able to produce retinotopic maps that resembled the quality of the standard pRF mapping both with and without eye movements.

Highly unusual for functional brain mapping studies of visual cortex, two of our participants in this experiment had an uncorrected refractive error. We deliberately did not correct their vision with spectacle lenses because that makes the eye tracking unreliable. However, this also enabled us to compare retinotopic maps and parameter estimates for the two analysis methods in the presence of optical blur. The pRF maps for a highly myopic participant (P3, refractive error: left = -3.5, right = - 4.25), contained considerable data loss even in the stable fixation condition (Supplementary Figure S2). There was a widespread ipsilateral pRF locations throughout the foveal and parafoveal representations of early visual cortex and the lateral occipital cortex. Especially in the early regions, these must constitute artifactual estimates. In the presence of large, random eye-movements, no map architecture could be distinguished at all. In contrast, CF maps revealed the expected pattern of polar angle reversals in occipital cortex, although the extent of activation was reduced compared to the other participants. Large eye-movements further reduced the map coverage somewhat, but the retinotopic architecture remained robust.

### Laser Kiwi experiment

Having shown that CF analysis is more robust to large eye movements, we then tested whether our method could generate retinotopic maps without controlled mapping stimuli and in the absence of any systematic fixation. This should further extend maps to peripheral locations: as the gaze shifts to the edge of the screen or beyond, the dynamic visual stimulus is effectively moved into the periphery. To this end, we specifically designed a video game called *Laser Kiwi* to induce large and frequent eye movements, covering the whole screen. We then generated CF maps from fMRI data acquired during this task. Note that it is impossible to analyze this experiment with the conventional pRF method.

The retinotopic maps obtained contained the expected retinotopic organization similar to conventional mapping experiments, at least in the early regions V1-V3 (Figure 6A). Moreover, despite some variability in estimates in ventral occipital cortex, in most participants the polar angle reversal corresponding to the lower vertical meridian was visible in V4, indicating its anterior border. V3A/B abuts the V3 dorsal quarter-field (V3d) and contains a full hemifield representation. Polar angle representations of V3A/B were consistent with previous findings [Larsson and Heeger, 2006; Wandell et al., 2007]. Further dorsal to these regions are retinotopic regions running across the medial wall of the intraparietal sulcus, comprising a series of foveal representations [Swisher et al., 2007]. We however only observed hints of the most posterior of these regions in our CF maps.

**Figure 6.**
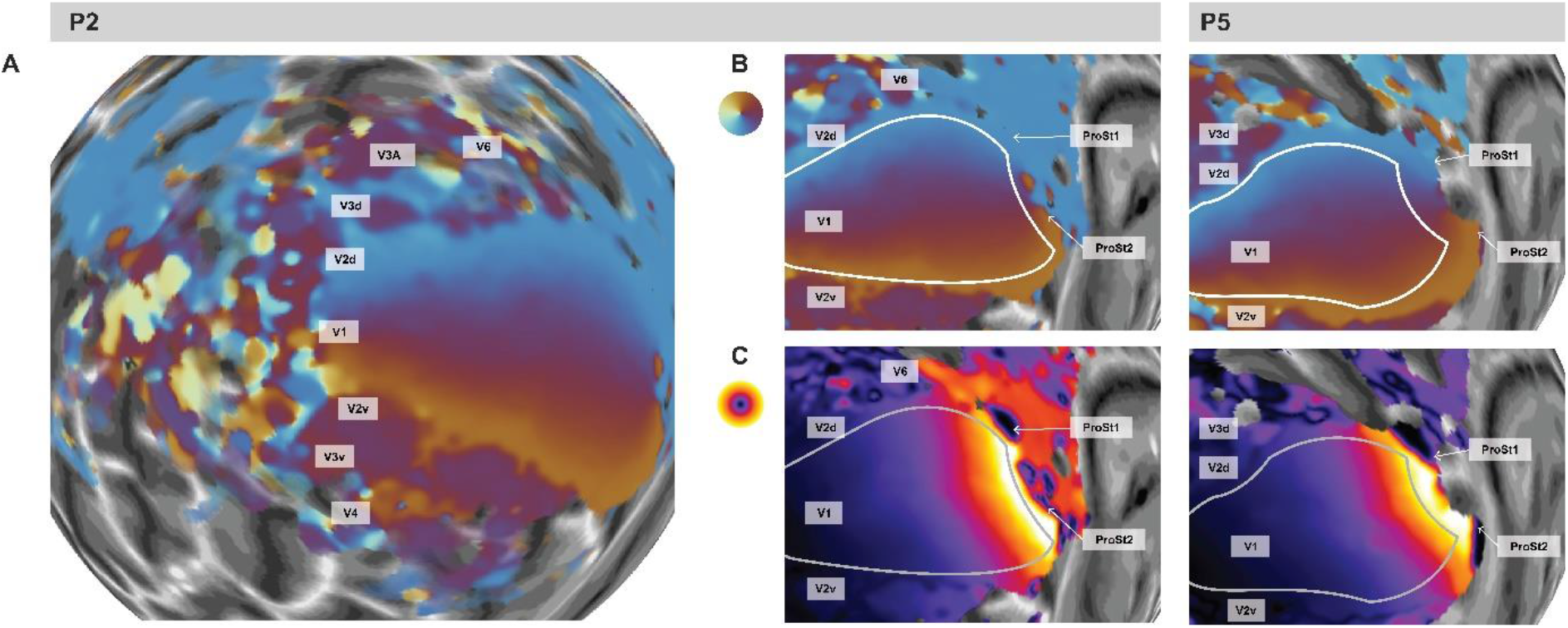
Retinotopic maps in the *Laser Kiwi* experiment. **A**. Polar angle maps derived with CF analysis of participant P2 shown on a spherical, inflated model of the left occipital cortex. **B**,**C**. Zoomed-in view of the representation of the peripheral visual field in early visual cortex in participants P2 and P5 (see grey bars). The outline indicates the borders of the template region V1. Maps show polar angle (**B**) and eccentricity (**C**).

MT+ is a motion sensitive cluster of areas situated on the border between lateral occipital and temporal cortex. This region is also known as the temporal occipital regions, comprising two full hemifield maps in TO1 and TO2 [Amano et al., 2009], separated by an upper vertical meridian representation that should merge in a foveal representation. Our maps for MT+ were ill-defined. Both polar and eccentricity maps were noisy, and relatively inconsistent across participants. However, a proportion of our MT+ maps represented the ipsilateral visual field, consistent with previous reports [Amano et al., 2009].

Area ProSt is located anterior to V1 in the fundus of the calcarine fissure and dominated by the visual periphery [Mikellidou et al., 2017; Yu et al., 2012]. It shares one of its borders with the V1 medial border representing an eccentricity reversal [Mikellidou et al., 2017]. Our CF maps were able to reveal substantial activation in ProSt. Across all participants, the ProSt polar angle gradient was like that of V1 (Figure 6B). The upper vertical meridian was represented at the ventro-medial border of the ProSt and continued to the lower vertical meridian on the dorsal side.

As expected, a considerable proportion of these regions represented the far periphery, extending up to 70-80° eccentricity. This peripheral coverage is farther than what was previously found with pRF mapping [Mikellidou et al., 2017]. The eccentricity map also exhibited a drastic reversal, presumably demarcating the border between V1 and ProSt. This reversal corresponded well with the anterior edge of the template region V1 defined by the probabilistic prediction [Benson et al., 2012]. Consistent with the atlas definitions [Sereno et al., 2022], we could identify two separate eccentricity reversals possibly corresponding to two ProSt regions, a dorsal region located in the cuneus and a ventral region on the lingual gyrus. Finally, CF sizes were also extremely large in these regions, both in ProSt and V1.

Area V6 is located anterior to V3d and parallel to V2d [Galletti et al., 1999; Pitzalis et al., 2015]. This area contains a full hemifield representation and, like ProSt, it is biased towards representing the visual periphery [Pitzalis et al., 2006]. CF maps of V6 were somewhat variable and inconsistent between participants (Figure 6B), although some polar angle gradients running orthogonal to those of V3d and V2d, and representing more than the upper quadrant, could be distinguished. Some eccentricity maps also contained a distinct foveal representation that might correspond to V6.

### Comparing CF and pRF sizes between tasks

Finally, we quantified CF sizes obtained from both the *Unstable Eye* and *Laser Kiwi* experiments (Figure 7A). Average CF sizes increased somewhat along the visual processing hierarchy and were considerably larger in V6 and ProSt. Interestingly, CF sizes were comparable in both experiments irrespective of eye-movements, except for slightly larger CFs in V3 in the *Laser Kiwi* experiment.

**Figure 7.**
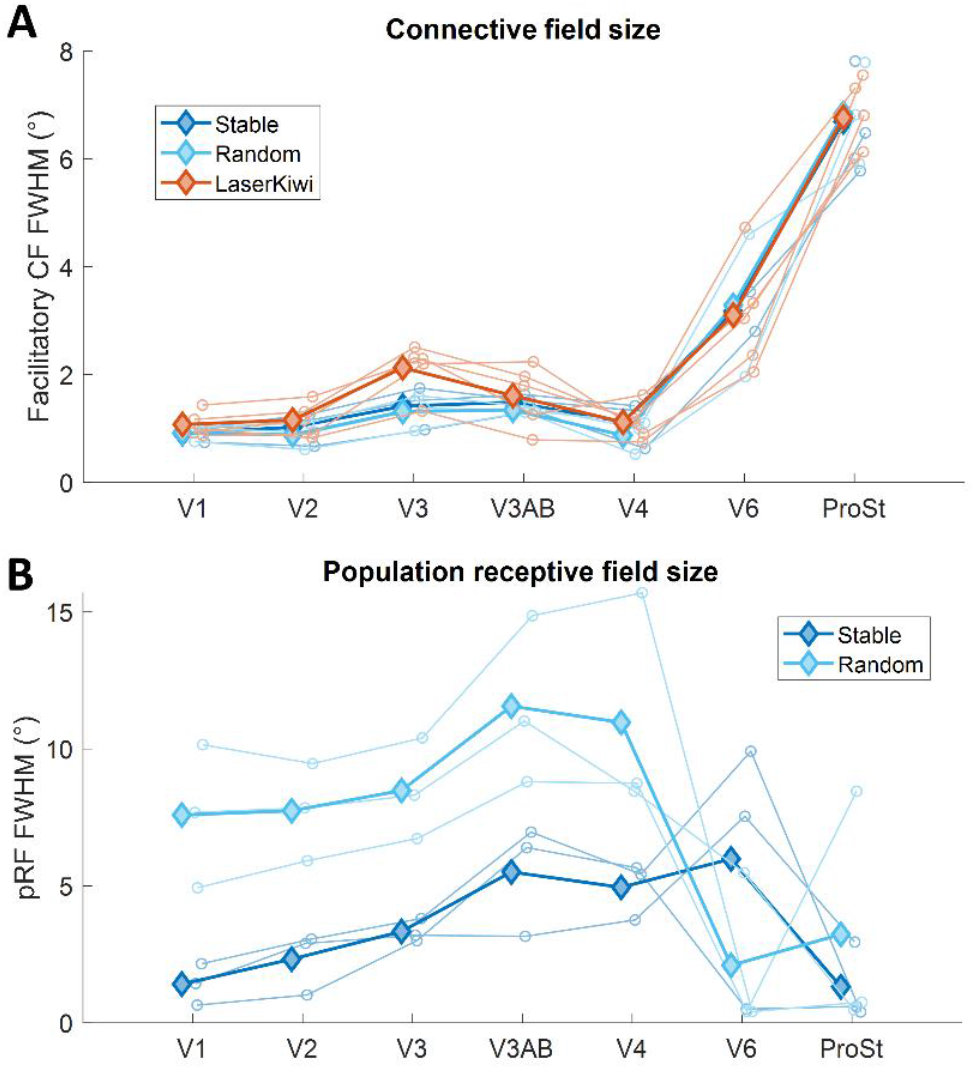
Comparison of mean CF (**A**) and pRF (**B**) sizes in different visual regions. Solid lines and filled diamonds show the mean across participants. Dotted lines and unfilled circles denote data from individual participants. Blue curves show data from the *Unstable Eye* experiment: Dark blue: stable fixation condition. Light blue: random fixation condition. The red curve in A shows data from the *Laser Kiwi* experiment.

For comparison, we also investigated changes in pRF size in the *Unstable Eye* experiment (Figure 7B). Consistent with CF sizes, average pRF sizes increased along the visual hierarchy. However, there were a few noticeable differences between CF and pRF size. First, even with stable fixation pRFs were overall much larger than CFs. This is consistent with our analysis of the *Standard Mapping* experiment (Figure 3). The averaged pRF sizes for visual regions V1-V3A/B ranged from 0.6° to approximately 7.0° while CF sizes contained a much smaller range from approximately 0.8° to 2.5°. Second, pRFs were unsurprisingly much larger in the presence of eye-movements owing to the greater variability of stimulated visual field locations than is captured by the pRF model. Third, while CFs were larger in peripherally biased area V6 and ProSt than in earlier regions, pRFs were notably smaller in those regions. Probably this is because only very few vertices contained significant pRF fits and these data are largely artifactual.

## Discussion

Here we used connective field analysis based on reversed correlation and a probabilistic atlas to map the retinotopic organization of human visual cortex, both under standard steady-fixation conditions and with unconstrained eye movements. Maps derived with standard retinotopic mapping stimuli showed that CF analysis is capable of reproducing retinotopic maps of similar quality as typically obtained with pRF analysis. This replicated previous work showing that we can predict response properties in distinct parts of the brain, allowing us to reconstruct the visual functional architecture [Haak et al., 2012; Knapen, 2021]. However, unlike those previous studies, we did not use empirical retinotopic maps for the template region V1 but instead capitalized on reports that cortical folding alone can reliably predict retinotopic maps in V1 [Benson et al., 2012]. This demonstrates that retinotopic mapping analysis is unnecessary for obtaining maps using the CF method. While the probabilistic atlas approach may contain idiosyncratic errors, it is sufficient for revealing retinotopic organization. Our CF maps also covered a greater amount of the cortex, both by revealing map structure outside the directly stimulated visual field and generally less patchy maps. The neural-referred nature of the CF analysis means that the mapping of each voxel in the brain does not require the systematic modelling of stimulus location and timing within the visual field.

We then tested an assumption about the CF method that follows from its independence of systematic stimuli: it should also be possible to obtain retinotopic maps with the CF method under unconstrained eye movements. Conventional retinotopic mapping (such as with the pRF method) necessitates steady fixation. Even though eye-tracking can be used to correct the stimulus model to improve pRF mapping results [Hummer et al., 2016], the corrections can only compensate for small eye-movements. Eye-movements will therefore still degrade retinotopic map quality. In contrast, the CF method does not “correct” for eye-movements; rather, it is robust to the deleterious effect of eye-movements because the analysis is relative to cortico-cortical connections instead of relying on a model of the visual stimulus in the first place. As predicted, despite considerable eye-movements CF maps were accurate and reliably revealed the retinotopic organization as expected from maps obtained with stable fixation. Meanwhile, maps obtained with pRF analysis were severely impacted by eye-movements.

Eye-movements also increased estimates of pRF size, presumably because unstable fixation results in greater variance of stimulated visual field locations. We found no such influence of eye-movements on estimates of CF sizes, again illustrating that the neural-referred nature of CF analysis obviates the need for stable fixation.

Optical defocus and a blurred retinal image, such as caused by myopia, can also impact the quality of retinotopic maps. In theory, the same reason why CF is robust to eye-movements predicts it should also be more robust to image blur. We did not specifically set out to manipulate image blur in the present study. However, we included participants with uncorrected refractive error out of necessity: our experiments required eye tracking which is impossible when the participant wears corrective spectacles. In one participant with significant myopic defocus (left = -3.5, right = -4.25), we indeed observed considerably deterioration of pRF maps, especially within foveal and parafoveal representations. In contrast, maps obtained with CF analysis were much more complete, and revealed the retinotopic architecture. We note, however, that a blurred retinal image likely increases the spatial correlations between voxels in the template region V1 and therefore still exerts some effect on CF estimates.

By removing the need for focused and systematic stimuli, CF mapping should generally be more versatile. Previous work has shown that CF maps can be estimated even with resting state data when the participant’s eyes are closed or while they watch movies [Gravel et al., 2014; Knapen, 2021]. Using ecologically valid and engaging stimuli should improve general task compliance and also make retinotopic mapping more suitable for use in patients and subject populations that struggle with the constraints of standard methods (e.g., participants at either end of the life span, individuals with nystagmus, or other non-neurotypical participants). However, those previous studies only showed some map structure in early visual cortex; one of them primarily discussed changes in retinotopic eccentricity tuning under different task conditions [Knapen, 2021]. We reasoned that by using an engaging task that encourages large eye-movements, we could widen the field of view being stimulated and thus enhance the mapping of the peripheral visual field: fixations at the screen edge move most of the screen into the periphery and therefore evoke visually-driven signals corresponding to those locations.

We therefore designed a gaze-contingent video game where participants shot at targets by fixating them. CF analysing of this data yielded retinotopic maps beyond V1-V4, including some coverage of regions known to be particularly dominated by the peripheral visual field, like ProSt and V6. We revealed a continuous polar angle map across the border between V1 and ProSt. This stands in contrast with a previous fMRI study that showed a polar angle gradient in ProSt running orthogonal to the one in V1 [Mikellidou et al., 2017]; however, the polar angle maps in that study were highly variable between participants and potentially limited by the scanning resolution. Importantly, a continuous polar angle maps between V1 and ProSt conforms with electrophysiological mapping of ProSt in the marmoset [Yu et al., 2012]. Generally, in earlier work using standard methods, the majority of eccentricity estimates fell between 45-60° [Elshout et al., 2018; Mikellidou et al., 2017; Stenbacka and Vanni, 2007; Yu et al., 2012]. For example, one study attempted to delineate peripheral visual field representation by using multi-focal stimuli together with optical near-view system [Stenbacka and Vanni, 2007]. They could stimulate eccentricities up to approximately 50° allowing them to locate V6. Delineation of this region was difficult, however, as responses could not be clearly distinguished from those of V2d and V3d. Our CF maps meanwhile contained the expected organization of V6. Similarly, in previous studies that utilized wide-view presentation systems to measure the peripheral representation, the maximum stimulation was limited to approximately 60° eccentricity [Mikellidou et al., 2017]. In contrast, our CF maps allowed us to obtain eccentricity estimates beyond 70°, without the need for continuous fixation, checkerboard stimuli, and unusual viewing conditions, thus enhancing participant comfort.

Consistent with previous reports for receptive field size [Mikellidou et al., 2017; Yu et al., 2012], we also found that CF sizes in ProSt were huge, spanning a considerable proportion of the visual field. We generally found no effect of the different tasks (stable, random eye-movements in the *Unstable Eye* experiment or unconstrained eye movements in the *Laser Kiwi* experiment). Thus, the large CFs in ProSt and the other representations of the periphery must reflect the underlying processing rather than eye-movement artifacts. However, we did not design our experiments to reveal subtle changes in CF size between conditions. As CF maps can change considerable under different task conditions [Knapen, 2021] it is possible that CF sizes will also change. Detecting this will require large samples with a within-subject design. CF sizes could change depending on the focus of spatial attention to local vs global stimulus attributes, or due to perceptual modulation by contextual visual illusions. A recent study investigating an illusory disappearance of visual stimuli (“artificial scotoma”) demonstrated considerable shifts in CFs depending on the participant’s perceptual state [Carvalho et al., 2021].

Naturally, even though retinotopic maps obtained with the CF and pRF methods are very similar, this does not imply they both measure the same underlying properties of brain function. At the coarse scale, pRF and CF sizes both increased as a function of eccentricity and when moving up the visual processing hierarchy. However, the pattern of CF sizes predicted pRF sizes only modestly in most regions, and were effectively uncorrelated in V1 and V2. The visual region corresponding to the CF is not the same as the pRF. While the latter describes how a voxel responds to stimulation across a range of visual field locations, the CF denotes the coactivation of neighboring voxels. As such, CF sizes cannot be interpreted as accurate measurements of pRF properties. Such estimates require estimating how responses vary when stimulating different visual field locations. Rather, CF size is probably related to cortical magnification. In line with that, we also estimated the extent of the inhibitory CF, characterizing neuronal populations in the template region that were inversely correlated with the target voxel. This inhibitory subfield was particularly large for foveal and parafoveal CFs, but exponentially reduced in size as a function of eccentricity. Such a pattern is to be expected if the inhibitory CF subfield is linked to cortical magnification and implies that the cortical extent of this inhibition is constant across V1.

Our CF method provides a promising alternative for mapping of human visual cortex, especially in participants where traditional methods are difficult to use. However, there are several aspects that could be improved. While the probabilistic atlas used for the template region was effective, slight discrepancies in functional organization between the atlas and the idiosyncrasies of the individual retinotopic architecture could result in some errors and misclassification of visual regions. Better atlases procedures improve the accuracy of the maps, such as predictions based on deep convolutional networks [Ribeiro et al., 2021]. Moreover, we also did not extensively filter our time courses. Many functional connectivity studies, especially for resting state analysis, use bandpass filtering to restrict the signal to low frequency fluctuations. Filtering can also remove of other nuisance factors, such as respiration, heartbeat, and signals from different brain tissues by using independent component analysis.

To our knowledge, CF analyses have exclusively used V1 as a template region. Future vision research could adopt different template regions, such as the frontal eye fields to investigate the cortico-cortical mapping of connections during attentional allocation, saccadic eye movements, and visual awareness. Moreover, CF analysis could of course also examine maps for other sensory modalities, such as tonotopic or somatosensory mapping. However, it is crucial to use accurate template maps as a basis for the template region. Unless empirical maps are used, this entails the creation of atlas-based maps for auditory and somatosensory cortex. Either way, the approach we present here using CF, engaging stimulus protocols, and unconstrained fixation opens new opportunities for studying visual field maps in the human brain.

## Acknowledgments

We thank the staff at CAMRI for their help with data collection. This research was supported by a CAMRI pilot grant and a Research Development Fund award to DSS.

## Data availability

For the *Unstable Eye* and *Laser Kiwi* experiments processed map data are publicly available at https://zenodo.org/record/7117929. Analysis and stimulus presentation code is available at https://osf.io/46wcm. National data protection rules prohibit the sharing of potentially identifiable brain images. Conditional sharing of raw data may be arranged with the corresponding author upon request.

## Supplementary Information

**Supplementary Figure S1.**
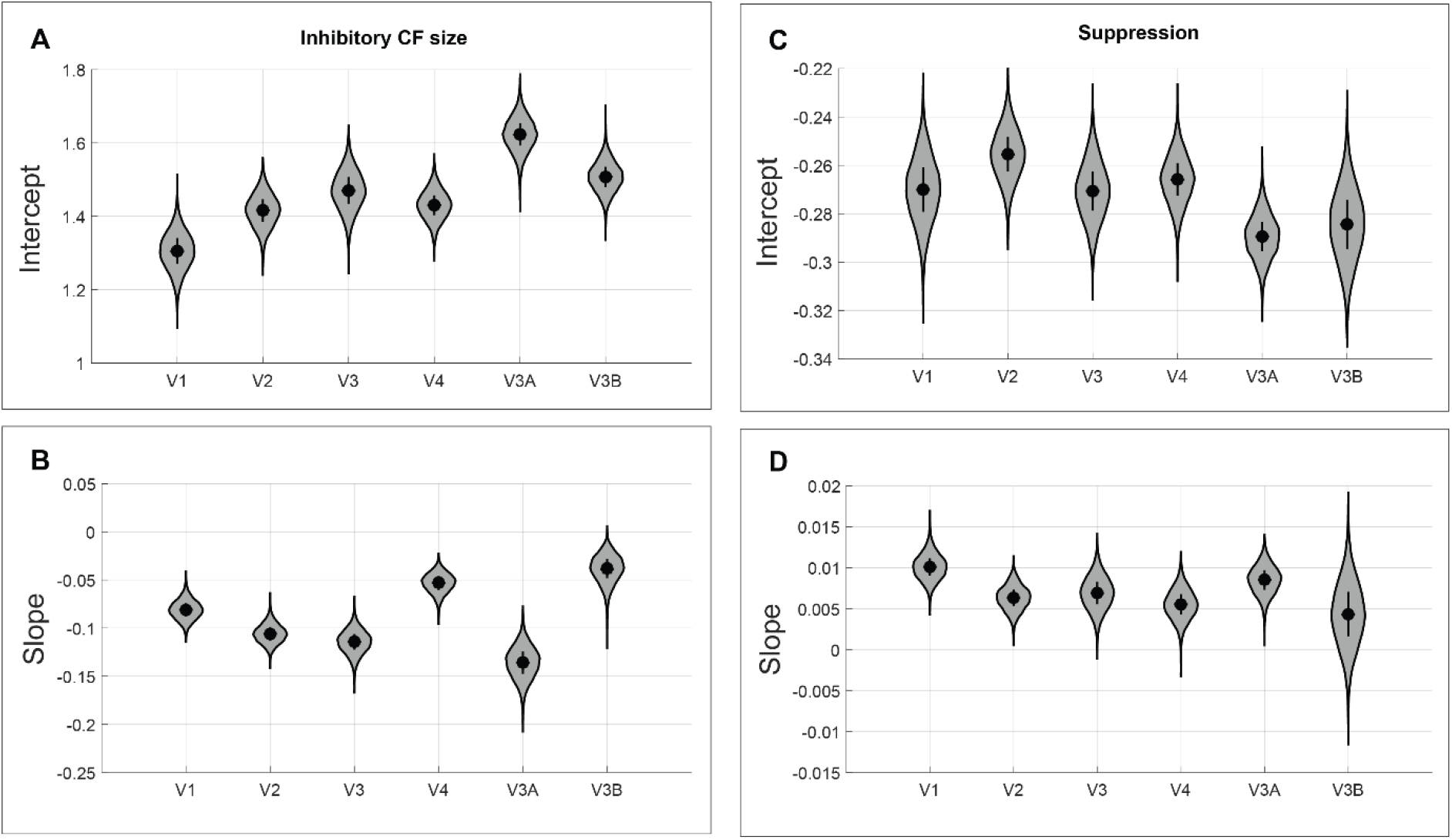
The panels show the mean intercept (**A**,**C)** and slope **(B**,**D)** coefficients from linear regression between eccentricities and inhibitory CF size (**A**,**B**) and suppression strength (**C**,**D**), respectively. Symbols and error bars denote the group mean parameter ±1 standard error. Violin plots show the bootstrapped distribution smoothed with a kernel function. For inhibitory CF size, data were log-transformed before linear regression.

**Supplementary Figure S2.**
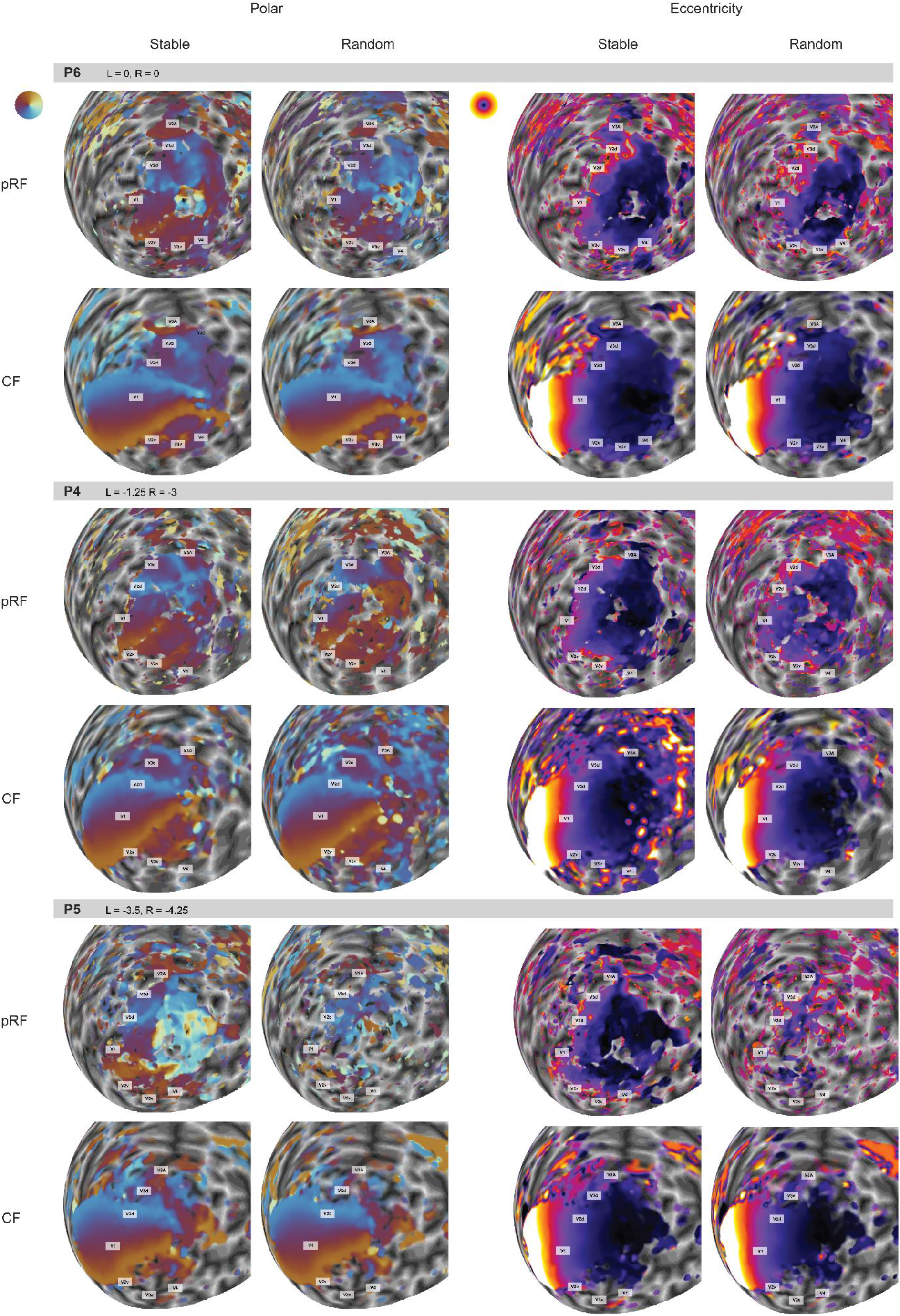
The effect of large eye movements on retinotopic maps in the *Unstable Eye* experiment of all subjects participated in the task (P4-P6). Polar angle (left) and eccentricity maps (right) were derived by pRF and CF analysis (rows), either with stable or random fixation patterns (columns). Each panel shows maps of given participant. The gray bars denote the participant identified and their refractive error in the left (L) and right (R) eyes.

